# Impact of dendritic spine loss on excitability of hippocampal CA1 pyramidal neurons: a computational study of early Alzheimer disease

**DOI:** 10.1101/2024.01.20.576500

**Authors:** Chengju Tian, Isabel Reyes, Arjun V. Masurkar

## Abstract

Synaptic spine loss is an early pathophysiologic hallmark of Alzheimer disease (AD) that precedes overt loss of dendritic architecture and frank neurodegeneration. While spine loss signifies a decreased engagement of postsynaptic neurons by presynaptic targets, the degree to which loss of spines and their passive components impacts the excitability of postsynaptic neurons and responses to surviving synaptic inputs is unclear. Using passive multicompartmental models of CA1 pyramidal neurons (PNs), implicated in early AD, we find that spine loss alone drives a boosting of remaining inputs to their proximal and distal dendrites, targeted by CA3 and entorhinal cortex (EC), respectively. This boosting effect is higher in distal versus proximal dendrites and can be mediated by spine loss restricted to the distal compartment, enough to impact synaptic input integration and somatodendritic backpropagation. This has particular relevance to very early stages of AD in which pathophysiology extends from EC to CA1.

## Introduction

Information processing by neurons relies on robust integration of synaptic input which ultimately leads to action potential initiation, the principal currency of information flow through neural circuits. Disruption of this process in disease states may underlie cognitive and behavior symptoms. For example, synaptic degeneration is a hallmark of Alzheimer disease (AD) (Terry et al., 1991; Sze et al., 1997, Scheff et al 2007), thought to be driven by amyloid and tau pathology and which may precede frank neurodegeneration (Tzioras et al., 2023). Specifically, area CA1 of hippocampus is implicated in early stages with respect to AD pathology and significant synaptic loss, which may contribute to amnestic deficits in pre-dementia stages (Braak and Braak, 1991; Scheff et al., 2007). According to this pathophysiologic model, CA1 pyramidal neurons (PNs) at this stage may show decreased action potential output due to reduced synaptic drive.

An alternative impact of spine loss is suggested by a study that linked compromised dendritic architecture of CA1 PNs in an AD transgenic model to increased neuronal excitability (Siskova et al., 2014). The authors demonstrated that loss of dendritic membrane, and subsequently of dendritic leak currents, results in an increase of overall input resistance and consequently an aberrant amplification of remaining synaptic inputs. This may contribute to the hyperexcitability mechanisms thought to impact cognitive symptoms and AD pathophysiology (Palop et al., 2007; Cirrito et al., 2005; de Calignon et al., 2012; Liu et al., 2012).

This leads to critical questions that pertain to even earlier stages of structural compromise in which spine loss occurs in absence of dendritic degeneration (Hsieh et al., 2006; Bittner et al., 2010). Does spine loss alone, with intact dendrites, lead to increased excitability in CA1 PNs? Additionally, the CA1 apical dendrite is compartmentalized, with the proximal dendrite receiving CA3 input and the distal dendrite receiving input from entorhinal cortex, which is implicated in very early stages of AD (Braak and Braak, 1991; Braak and Braak, 1997; Lace et al., 2009). Thus, can spine loss limited to the much smaller distal apical dendrite alone have an impact on excitability? Given the heterogeneity of CA1 PNs according to somatic depth (Masurkar, 2018), are superficial (sPN) versus deep PNs (dPN) differentially impacted by these changes? And lastly, does spine loss have other impact on information processing by CA1 PNs? Here leverage passive, multicompartmental computational models of CA1 PNs to simulate spine loss and assess its effects to answer these questions.

## Methods

CA PN models were created from morphologies previously derived from whole cell slice recordings and detailed in a previous study (Masurkar et al., 2020). Morphometric analysis was performed using the ImageJ Sholl analysis plugin (Ferreira et al., 2014), which was subsequently utilized to compute the Branching Index (Garcia-Segura and Perez-Marquez, 2014):

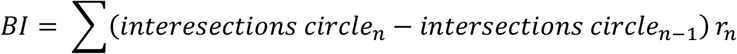

The NEURON modeling suite (Hines and Carnevale, 1997) was used to create passive, multicompartment models of 4 sPNs and 4 dPNs and perform simulation experiments. As detailed in our prior study (Masurkar et al., 2020), parameters fit included R_in_ (sPN: 151 MΩ; dPN: 207 MΩ; achieved by adjusting g_pas_), R_a_ = 35.4 Ω-cm, C_m_= 1 μf/cm2, e_pas_= -70 mV. The The method to model spines was also as previously described (Srinivas et al., 2017; Masurkar et al., 2017). We recapitulate this method here. In brief, we used the same spine densities (spines/μm) as derived in our prior work (Masurkar et al., 2017; Masurkar et al., 2020): sPN: 1.09 in distal apical dendrite, 1.46 in the proximal apical dendrite; dPN: 0.54 in distal apical dendrite, 1.37 in the proximal apical dendrite. To correct for spines in the dendritic shaft plane, a correction factor was used (Feldman and Peters, 1979) using our measurements of spine density (Sd) and dendrite radius (Dr), and published (Harris and Stevens, 1989) values for spine length or Sl (0.7 μm) and spine head diameter or Sd (0.35 μm):

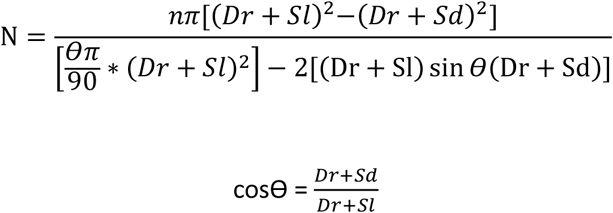

The “SpineScale” (Bush and Sejnowski, 1993; Stuart and Spruston, 1998; Golding et al., 2005; Routh et al., 2009) was used to calculate the extra membrane surface area contributed by spines for every 1μm of dendrite, in which d is dendrite branch diameter, N is corrected spine density, and SA is spine surface area (0.85μm^2^) based on previously published measures of head and neck area (Harris and Stevens, 1989):

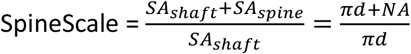

Dendrites were designated as distal or proximal dendrite based on the original morphology as visualized within the brain slice of origin. Spines in these dendritic compartments were subsequently modeled by a reduction in local resistance R_in_(computed as R_in_/SpineScale) and an increase in local capacitance C_m_(computed as C_m_*SpineScale). Consequently, a decrease in spines was modeled as an increase of R_in_and a decrease in C_m_. No other active conductances were added to the model.

Simulations were performed with a dt = 0.025. The AlphaSynapse point process (tau = 1 ms, g_max_ = 0.003 μS, e_rev_ = 0 mV) was used to simulate glutamatergic input at random locations in the proximal or distal apical dendritic tuft, with a recording electrode at the soma to record the EPSP in current clamp. Input resistance at the soma was measured using a 1500ms negative square pulse of current injection (I_input_ = -10 pA) and measuring ΔVm/I_input_. The Impedance function was used to plot dendrosomatic or somatodendritic attenuation as a function of frequency by choosing a representative dendritic location and the soma for recording or stimulation, as relevant. Statistical analysis was performed using Graphpad Prism 9. Means were compared using t-test, while graphs were compared using 2-way ANOVA.

## Results

We first asked if synaptic responses in the distal dendrite were differentially affected by overall spine loss compared to synaptic responses in the proximal dendrite. To answer this and subsequent questions in this study, we generated passive multicompartment models of 4 CA1 sPNs and 4 CA1 dPNs (Figure 1A, left), derived from Neurolucida tracing of morphologies visualized via biocytin staining (see Methods). The models incorporated contributions from experimentally-derived spine densities to dendritic resistance and capacitance. Recordings of the somatic excitatory postsynaptic potential (EPSP) were simulated in response to the stimulation of multiple (n = 20) synaptic locations in the distal dendrite or proximal dendrite, the sites of entorhinal cortex and CA3 inputs, respectively, with spine density in each compartment equivalently reduced by 20-80% (Figure 1A, right).

**Figure 1.**
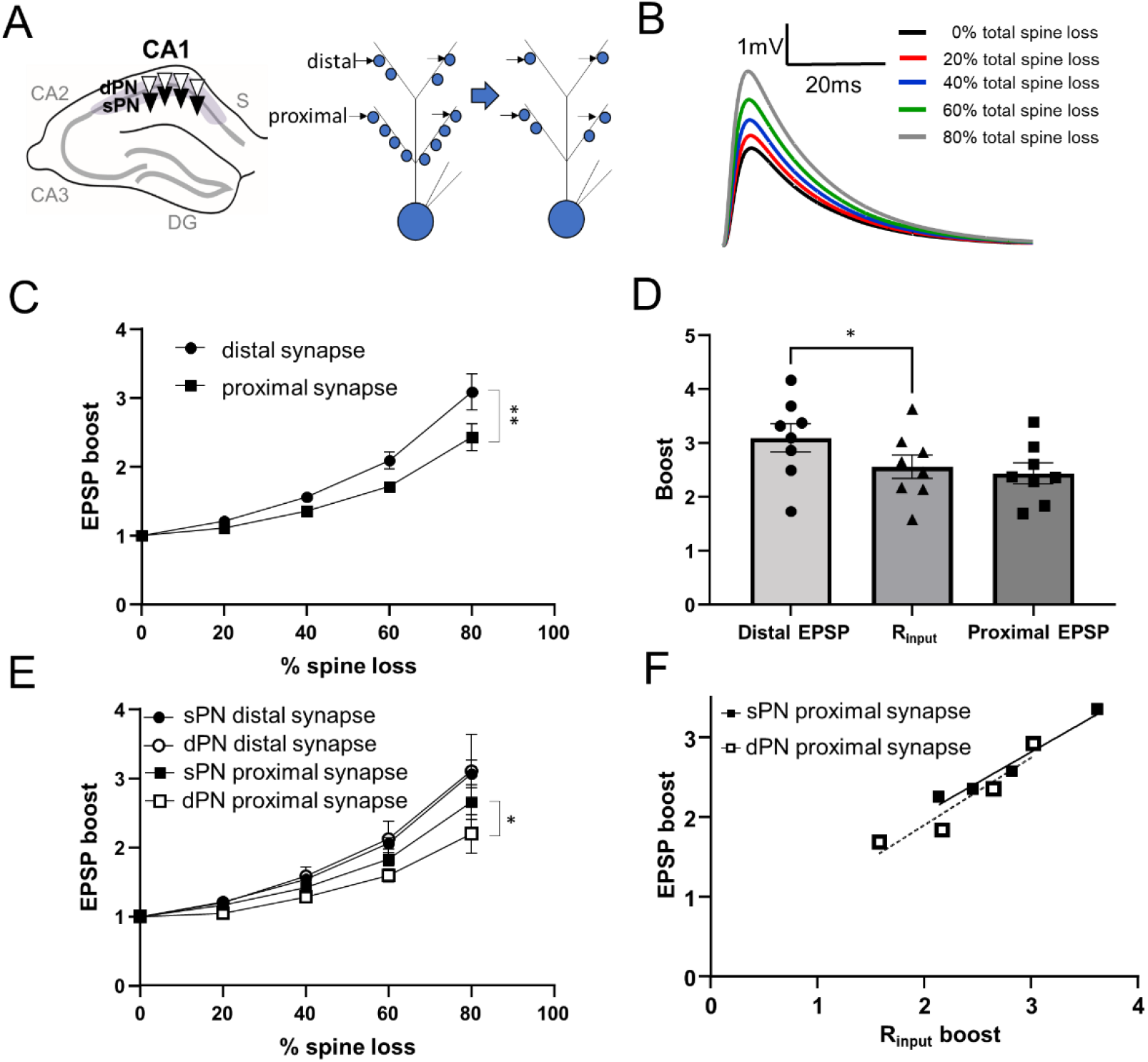
Simulation of the impact of spine loss across the apical dendrite on synaptic responses. **A**. Left: Schematic showing superficial and deep pyramidal neuron (sPN, dPN) layers in CA1, Right: Schematic of simulation in which synapses (n=20) are activated in distal or proximal apical dendrite compartments before and after varying degrees of uniform spine loss across both compartments. Excitatory postsynaptic potential (EPSP) is recorded at the soma. **B**. Overlay of traces from stimulating the distal dendrite of an sPN with % spine loss varied showing increased amplitude with increasing spine loss. **C**. Comparison of this EPSP boosting of distal and proximal synaptic sites as a function of % spine loss. **D**. Comparison of boosting in distal and proximal EPSP versus boosting of somatically-measured input resistance R_input_. **E**. Plots of distal and proximal EPS boosting vs. % spine loss in sPNs and dPNs. **F**. Plot of proximal EPSP boost vs. Rinput in sPNs (R^2^=0.9644, p=0.0180) and dPNs (R^2^=0.8960, p=0.0534). For all, n=8 (4sPNs, 4dPNs), *p<0.05, **p<0.001.

With increasing spine loss, somatically-recorded EPSPs increased in amplitude (Figure 1B). When plotted as a function of total apical spine loss, this “boost” of the EPSP (compared to no spine loss) was surprisingly higher for distal compared to proximal EPSPs (Figure 1C). During these simulations, we also measured somatic input resistance (R_input_) via a small negative current injection (see Methods). With a reduction in total spine density, somatic R_input_ increased to the same degree as the boost of the proximal EPSP, but was less than the boost of the distal EPSP (Figure 1D, 80% spine loss). Thus, the boost of the distal EPSP appeared to be additionally driven by local effects on dendritic excitability. We next examined if the EPSP boost from spine loss was differentially evident in superficial versus deep PNs (sPNs, dPNs). When dividing data by neuronal subpopulation, we found that there was no difference in EPSP boosting of distal inputs, but there was more boosting of the proximal dendrite EPSP in sPNs (Figure 1E). Given that the boost of the proximal EPSP is closely aligned with increases in R_input_, we surmised that this relationship may be stronger in sPNs, which have a shorter distance between their somata and the full extent of their proximal dendrite. Indeed, sPNs showed a statistically significant linear correlation between the increases in Rinput and the boost of their proximal dendrite EPSP at 80% spine loss, whereas this relationship was not statistically significant in dPNs (Figure 1F).

We then asked if the architecture of the dendritic arbor influences the degree to which spine loss leads to EPSP boosting. We focused on two features: branching complexity, given its influence on electrotonic properties (Rall, 1959), and dendritic length. We first assessed branching complexity (Figure 2A) via the Branching Index (Garcia-Segura and Perez-Marquez, 2014), derived from Sholl analysis (Sholl, 1953). For both proximal and distal EPSPs, there was no correlation between Branching Index and EPSP boosting (Figure 2BC). We next assessed if the relative length of the proximal dendrite (and its branches) influenced EPSP boosting (Figure 2D). In this case, the proximal EPSP boosting was significantly correlated with the percent of the apical dendrite comprised by the proximal compartment (Figure 2E), whereas the distal EPSP showed no such correlation (Figure 2F). The distal EPSP also did not show correlation with percent of dendrite comprised by the distal dendrite (not shown). This suggests that the boosting effect of the proximal EPSP is largely driven by spine loss in the proximal dendrite. Distal EPSP boosting may have more complex relationships to neuron morphology.

**Figure 2.**
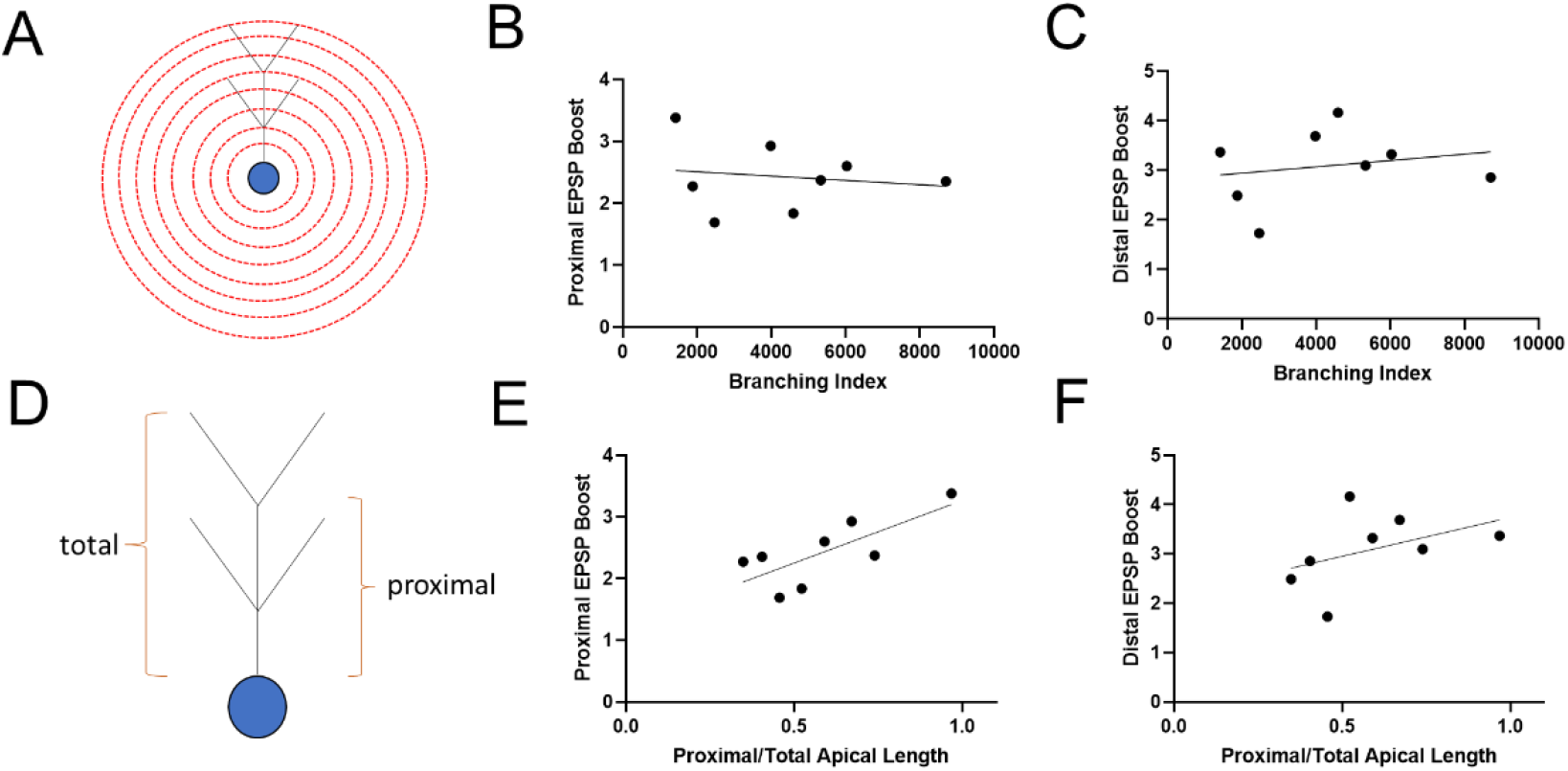
Morphological contributions to EPSP boosting due to spine loss. **A**. Schematic showing Sholl analysis used to then compute Branching Index **B**. Plot of proximal EPSP boosting versus Branching Index (R^2^=0.02357, p=0.5863). **C**. Plot of distal EPSP boosting versus Branching Index (R^2^=0.0463, p=0.6221). **D**. Schematic showing computing of proximal/total apical dendrite length. **E**. Plot of proximal EPSP boosting versus proximal/total apical length (R^2^=0.5586, p=0.0331). **F**. Plot of proximal EPSP boosting versus proximal/total apical length (R^2^=0.1827, p=0.2908). For all, n=8 (4 sPNs, 4 dPNs).

In the above simulations, the number of synapses stimulated was left unchanged. Yet with spine loss, there may be less synapses available to be stimulated. As such, we next asked how the EPSP boosting was affected if the number of stimulated synapses was scaled in accordance with the amount of spine loss (Figure 3A). Repeating our simulations with this condition in place, as expected the EPSP amplitudes were reduced with increasing spine loss (Figure 3B). However, when plotted against the expected degree of amplitude reduction (ex. 20% of control amplitude with 80% spine loss), the distal and proximal EPSPs both showed a more tempered reduction (Figure 3C). Moreover, the distal EPSPs showed less reduction compared to the proximal EPSPs. We subsequently divided the measured EPSP reduction by the expected reduction to reveal that the EPSP boost in this context increased with spine reduction, and more so for the distal EPSPs (Figure 3D). When we examined sPNs and dPNs separately, we again did not find any differences with distal EPSP boosting. However, contrary to when the number of synapses when left unchanged, with the number of stimulated synapses scaled to the amount of spine loss, there was no longer a statistically significant difference in the amount of boosting of the proximal EPSP between the two cell types (Figure 3E, F).

**Figure 3.**
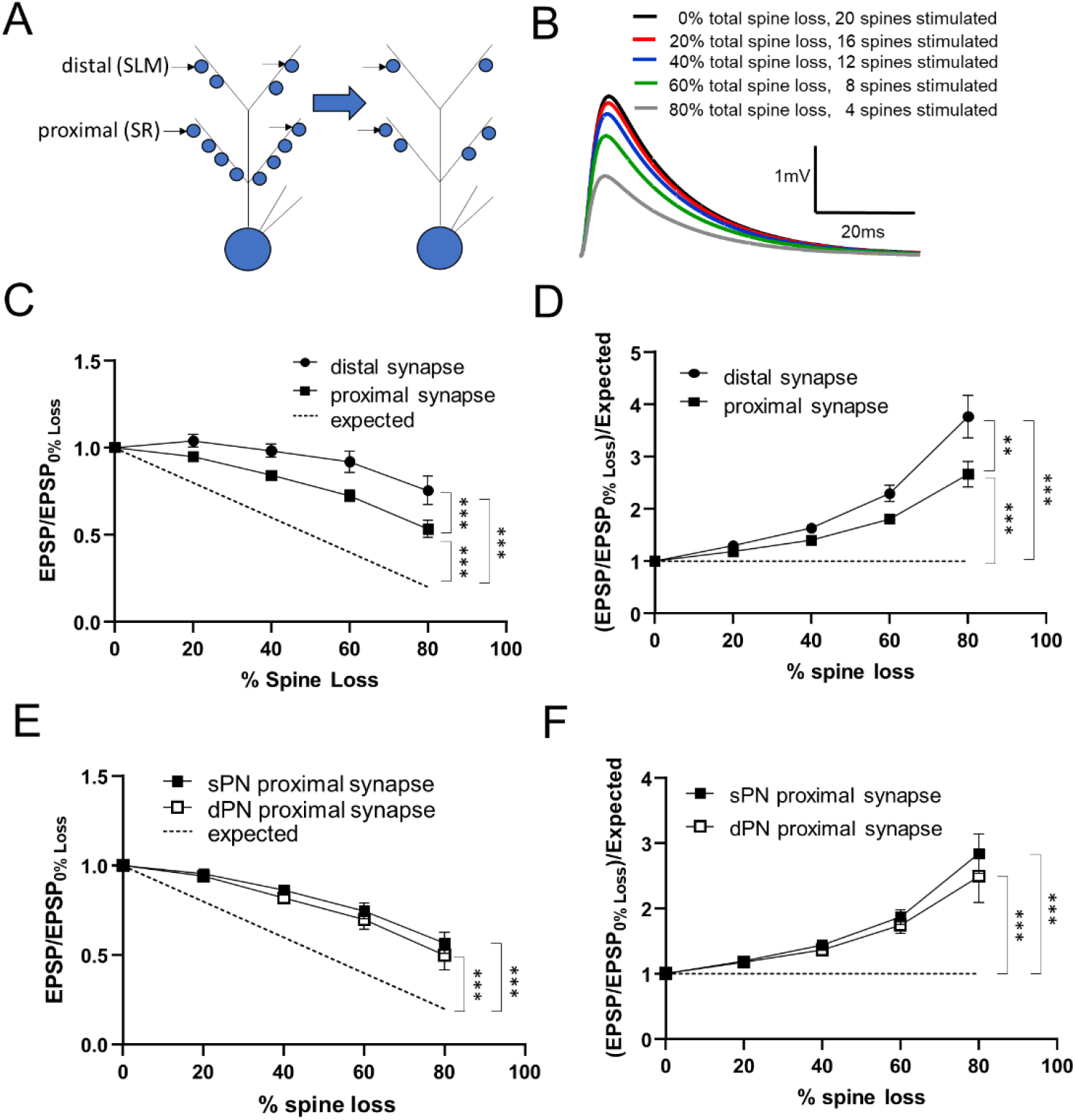
Impact of spine loss on synaptic responses with scaled activation of synapses. **A**.Schematic of simulation in which the number of synapses activated in the proximal and distal dendrite are scaled to the % spine loss. Excitatory postsynaptic potential (EPSP) is recorded at the soma. **B**. Overlay of traces from stimulating the distal dendrite of an sPN with % spine loss varied. **C**. Comparison of EPSP amplitude versus control (no spine loss, 20 synapses activated) of distal and proximal synaptic sites as a function of % spine loss. Dashed line represents expected relative reduction of EPSP amplitude without spine loss-mediated boosting. **D**. Boosting of distal and proximal EPSPs relative to expected reduction without spine loss-mediated boosting. **E**. Plots in C. separated between those from sPNs and dPNs. **F**. Plots in D. separated between those from sPNs and dPNs. For all, n=8 (4sPNs, 4dPNs), **p<0.0005, ***p<0.0001

We next asked to what extent spine loss-induced EPSP boosting could occur only in the presence of distal dendrite spine loss. We were motivated to study a scenario that would be more relevant to very early stage AD in which pathologic change begins to transition from EC to CA1 with concomitant changes in the distal dendrite (Braak and Braak 1991, Braak and Braak 1997). We first reduced distal dendrite spines by 20-80% and stimulated a constant number of synapses (n = 20) in the distal and proximal dendrite (Figure 4A). While the EPSP boost was notably less than when total spines were reduced, the distal EPSP still experienced a nearly 30% boost with 80% spine loss (Figure 4B). In contrast, the proximal EPSP showed very minimal change. We then performed these simulations with the number of stimulated synapses scaled to the degree of spine loss (Figure 4C). In this case, the distal synaptic response still slightly outperformed the expected decline (Figure 4D) and experienced greater than 40% boosting over the expected decline with 80% spine loss in the distal dendrite (Figure 4E). Similar to when total spine were reduced, there was no difference in this effect between sPNs and dPNs (not shown). While this comparative boosting appeared modest, during high frequency input bursts – those that can support long term potentiation – additive effects may amplify its impact. For example (Figure 4F), under these conditions a 100 Hz burst of 10 pulses (a component of theta burst stimulation) results in a compound synaptic potential approximately twice as large when the same number of synapses are activated with 80% spine loss compared to control conditions.

**Figure 4.**
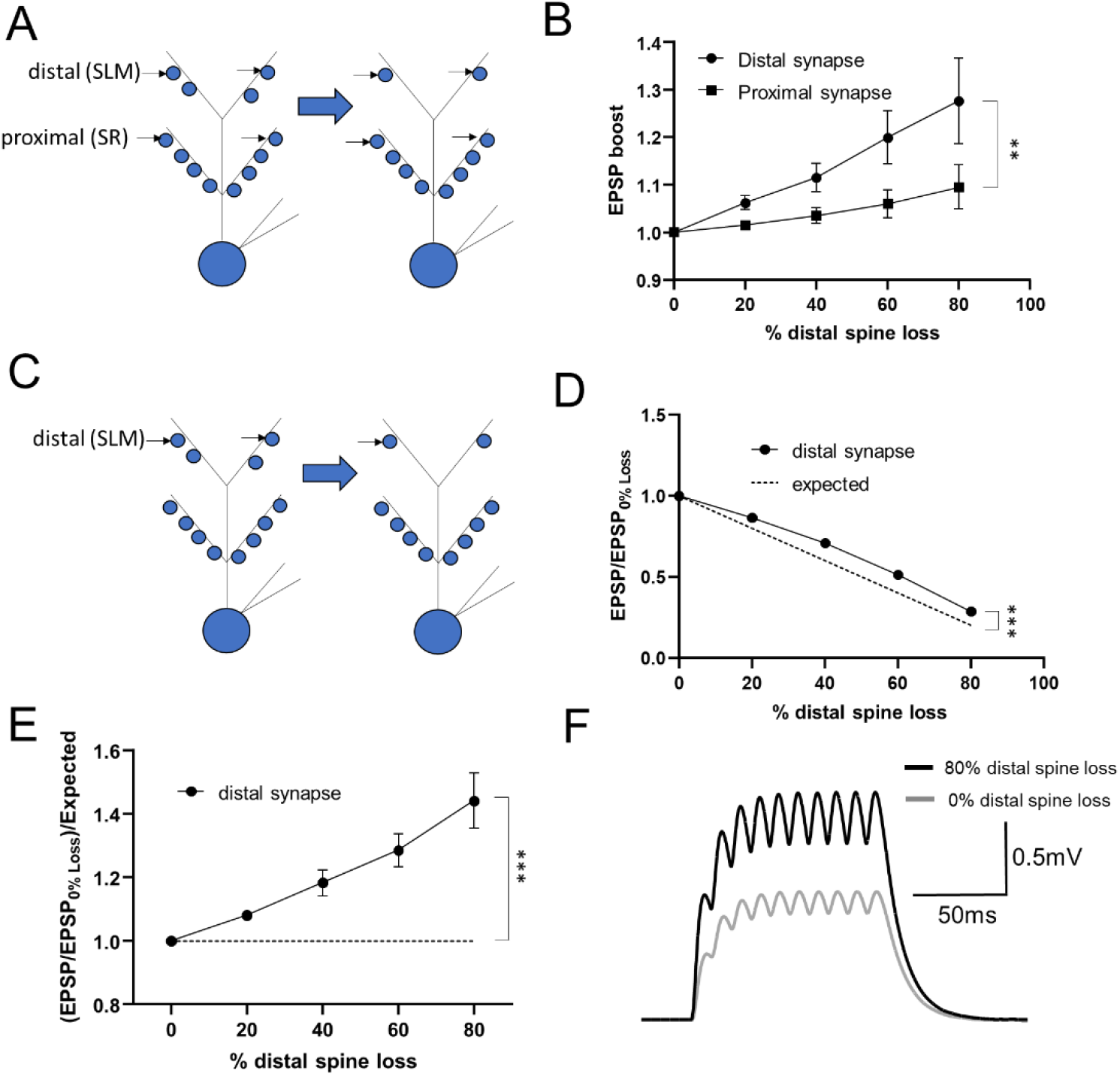
Impact of distal dendrite spine loss on synaptic responses. **A**. Schematic of simulation in which spine loss is limited to the distal dendrite and the number of synapses activated is constant (n=20). Excitatory postsynaptic potential (EPSP) is recorded at the soma.**B**.Degree of boosting of EPSP amplitude (compared to no spine loss) as a function of % spine loss, for distal and proximal dendrite synaptic activation. **C**. Schematic of simulation in which spine loss is limited to the distal dendrite and the number of synapses activated in the proximal and distal dendrite are scaled to the % spine loss. Excitatory postsynaptic potential (EPSP) is recorded at the soma. **D**. Comparison of distal EPSP amplitude compared to control (no spine loss, 20 synapses activated) as a function of % spine loss. Dashed line represents expected relative reduction of EPSP amplitude without spine loss-mediated boosting. **E**. Boosting of distal EPSPs relative to expected reduction without spine loss-mediated boosting. P<0.0001 **F**. Example trace of distal EPSP induced by stimulating 20 synapses with 10 x 100 Hz burst, with 80% (black) and without (gray) distal spine loss. For all, n=8 (4sPNs, 4dPNs), **p<0.001, ***p<0.0001

Since distal spine loss alone had an impact on distal EPSP amplitude boosting, we next asked if the loss of these spines impacted attenuation of such dendrosomatic propagation at different input frequencies. In this simulation (Figure 5A), voltage was injected at a distal dendritic site at various frequencies, and voltage response was measured at the soma. The degree of attenuation, V_soma_/V_dendrite_, was compared with intact distal dendrite spines versus with 80% loss of these spines (Figure 5B). In this context, while there was a mild improvement of attenuation across frequency with distal spines loss, it was not statistically significant, which may be in line with very local effects of distal spine loss. However, when we compared the degree of attenuation boosting, by dividing the two curves, across sPNs and dPNs, we noted that dPNs showed a greater degree of attenuation boosting that most notable at high frequencies (Figure 4C). We next simulated somatodendritic attenuation, which is relevant for action potential backpropagation and plasticity (Linden, 1999; Golding et al., 2001), in similar fashion (Figure 5D). In this context, there was less attenuation across all frequencies with distal spine loss (Figure 5E). In contrast to dendrosomatic attenuation boosting, there was no difference between sPNs and dPNs (Figure 5F).

**Figure 5.**
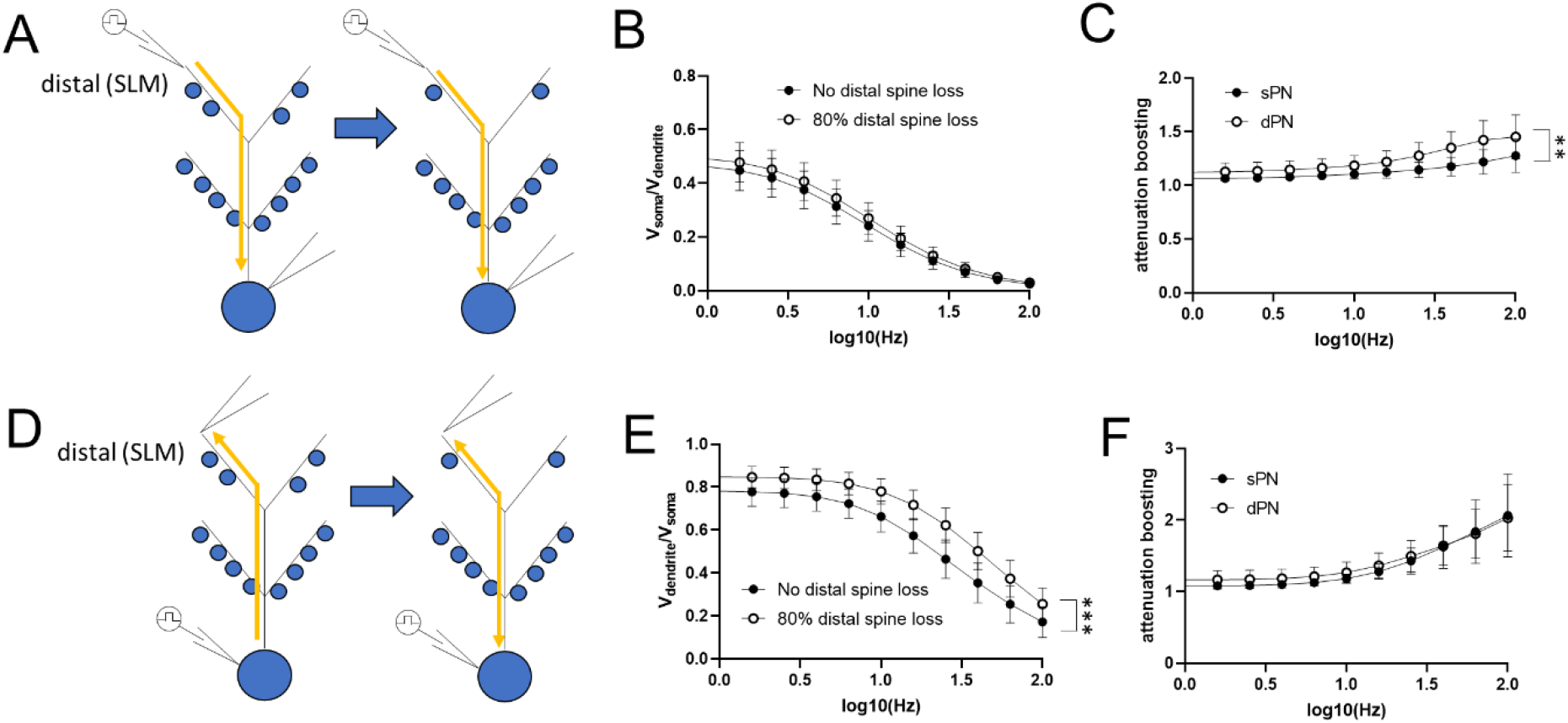
Impact of distal dendrite spine loss on dendrosomatic and somatodendritic propagation. **A**. Schematic of simulation in which spine loss (80%) is limited to the distal dendrite and the number of synapses activated is scaled to spine loss. To measured dendrosomatic attenuation, voltage input is given at the distal dendrite and voltage response is recorded at the soma. **B**. Plot of dendrosomatic attenuation versus frequency, with and without spine loss. **C**. Plot of degree of attenuation boosting (ratio of plots in B) in sPNs and dPNs. **D**. Schematic of simulation in which spine loss (80%) is limited to the distal dendrite and the number of synapses activated is scaled to spine loss. To measure somatodendritic attenuation, voltage input is given at the soma and voltage response is recorded at the distal dendrite. **E**. Plot of somatodendritic attenuation versus frequency, with and without spine loss. **F**. Plot of degree of attenuation boosting (ratio of plots in E) in sPNs and dPNs. For all, n=8 (4sPNs, 4dPNs), **p<0.01, ***p<0.0005

## Discussion

In this study, our simulations suggest that spine loss alone in CA1 PNs can mediate a boosting of the remaining inputs to their proximal and distal dendrites, targeted by CA3 and entorhinal cortex, respectively. Moreover, we show that this boosting effect is higher in distal versus proximal dendrites. Though the distal dendrite is a smaller compartment, this effect can also be mediated by loss in the distal dendrite alone to impact synaptic input integration and somatodendritic backpropagation.

In the context of AD, what may be the impact of such EPSP boosting? In early, preclinical stages, this phenomenon may serve as a compensatory mechanism to maintain synaptic drive of CA1 PNs. This may be especially relevant to the impact of the distal dendrite spine loss, as AD tauopathy begins in preclinical stages in entorhinal cortex and appears in CA1 in more prodromal stages (Braak and Braak, 1991; Bennett et al., 2005; Lace et al., 2009). As such, subtle dysfunction of this efferent area, as well as loss of related synapses in CA1, could be to some degree tolerated via this increased excitability. This boosting, in addition to improved somatodendritic attenuation to better permit action potential backpropagation, can maintain long term potentiation and optimal spatial coding that likely underlie memory-guided behavior (Linden, 1999; Hardie and Spruston, 2009; Bittner et al 2015). This is in line with other changes to spines that may have a similar protective role in early stages (Walker and Herskowitz, 2021). Alternatively, these changes may promote hyperexcitability, which appears to be an important early stage pathophysiologic mechanism that may even enhance spread of AD pathology (Palop et al., 2007; Cirrito et al., 2005; de Calignon et al., 2012; Liu et al., 2012). Given that hyperexcitability has been shown to arise in the entorhinal cortex (Xu et al., 2015; Rodriguez et al., 2020), compensation may have a maladaptive role by better facilitating propagation of this to CA1. It may have a multiplicative effect in turn via the enhanced intrinsic synaptic integration and capacity for action potential backpropagation, leading to maladaptive plasticity.

These effects are in line with other structural and physiologic changes in AD related to synaptic response and integration. Changes in spine morphology correlating with resilience have been noted (Boros et al., 2017, Boros et al., 2019) that may enhance synaptic current amplitude or flow through single spines and thus increase EPSP amplitude (Yuste, 2013). Voltage-gated ion channels particularly enriched in the distal dendrite, including the A-type potassium channel (Chen, 2005), HCN channel (Russo et al., 2021), and the GIRK channel (Martin-Belmonte et al., 2022), have all been noted to be decreased in AD models, which would potentiate our findings as well. What remains unclear is whether altered glutamate receptor expression within spines (Sze et al., 2001; Hsieh et al., 2006; Carter et al., 2004; Yeung et al., 2021) promote or temper these effects.

Given the increasing evidence that CA1 PNs are heterogeneous, including across its radial axis, we also examined if sPNs and dPNs were differentially susceptible to the impacts of spine loss. To this end we found that sPNs show more boosting of the unitary proximal dendrite EPSP (when proximal spines are also reduced), whereas dPNs show a greater enhancement of dendrosomatic propagation of distal input responses at high frequencies (when only distal spines are reduced). This may derive from baseline differences in morphology, with sPNs having a shorter distance between the soma and proximal dendrite. Furthermore, we selected sPNs and dPNs from a CA1 subregion in which sPNs, compared to dPNs, have a higher spine density in their distal dendrites with concomitant impact on distal EPSP kinetics (Masurkar et al., 2017; Masurkar et al., 2020). Thus, even with spine loss, the additional baseline capacitance from the extra spine density in sPNs may limit attenuation improvement at high frequencies compared to dPNs. Given that distal inputs are likely to be compromised prior to proximal input, given that entorhinal cortex is implicated in AD prior to CA3 (Braak and Braak, 1991; Lace et al., 2009), this would suggest that the dendrosomatic propagation changes (favorable to dPNs) would occur at an earlier stage than the enhancement of the proximal dendrite EPSP (favorable to sPNs). Given the distinct functional roles of sPNs/dPNs and distal/proximal inputs (Masurkar, 2018; Basu and Siegelbaum, 2016), how the evolution of these changes during AD impacts memory-guided behavior is an area of future work.

## Author contributions

AVM: conceptualization. CT, IR, AVM: performance of simulations. CT, IR, AVM: manuscript preparation.

## Funding

This work was funded by the BrightFocus Foundation (A2019602S), Alzheimer’s Association (AACF-17-524288), Blas Frangione Foundation, Leon Levy Foundation, and the National Institutes of Health (RF1AG072507, R21AG070880, P30AG066512).

## Competing Interests

None

## Notes

### Competing Interest Statement

The authors have declared no competing interest.

